## Supplementary figures and images for "Characterisation and chemometric evaluation of 17 elements in ten seaweed species from Greenland"

### Supplementary figure A

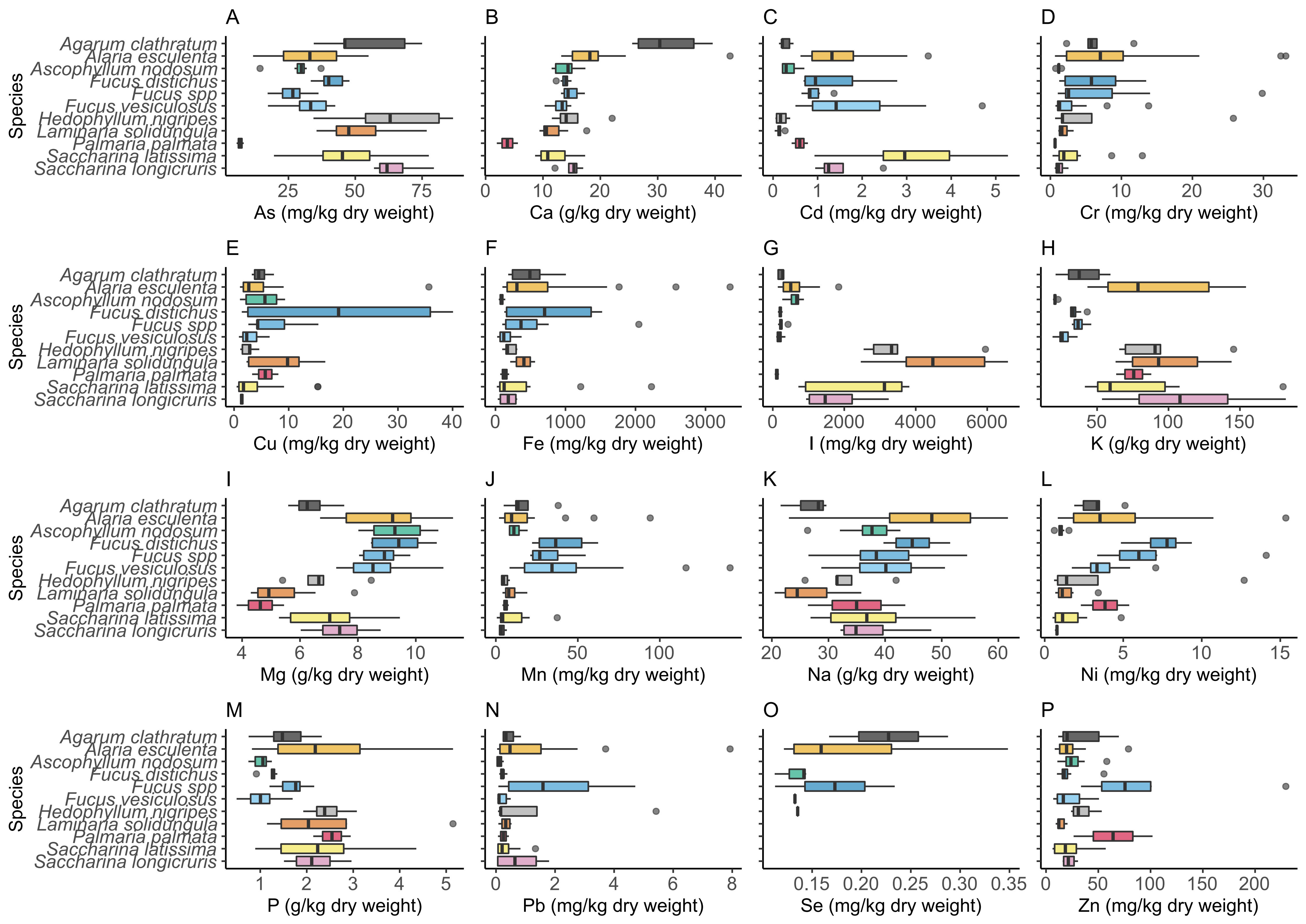
